# A chlorzoxazone-baclofen combination improves cerebellar impairment in spinocerebellar ataxia type 1

**DOI:** 10.1101/2020.08.14.251330

**Authors:** David D. Bushart, Haoran Huang, Luke J. Man, Logan M. Morrison, Vikram G. Shakkottai

## Abstract

**Background:** A combination of central muscle relaxants, chlorzoxazone and baclofen (chlorzoxazone-baclofen), has been proposed for treatment of cerebellar symptoms in human spinocerebellar ataxia (SCA). However, central muscle relaxants can worsen balance. The optimal dose for target engagement without toxicity remains unknown.

**Objectives:** Using the genetically precise *Atxn1*^*154Q/2Q*^ model of SCA1, we determine the role of cerebellar dysfunction in motor impairment. We also aim to identify appropriate concentrations of chlorzoxazone-baclofen needed for target engagement without toxicity to plan for human clinical trials.

**Methods:** We use patch clamp electrophysiology in acute cerebellar slices and immunostaining to identify the specific ion channels targeted by chlorzoxazone-baclofen. Behavioral assays for coordination and grip strength are used to determine specificity of chlorzoxazone-baclofen for improving cerebellar dysfunction without off-target effects in *Atxn1*^*154Q/2Q*^ mice.

**Results:** We identify irregular Purkinje neuron firing in association with reduced expression of the ion channels *Kcnma1* and *Cacna1g* in *Atxn1*^*154Q/2Q*^ mice. Using *in vitro* electrophysiology in brain slices, we identify concentrations of chlorzoxazone-baclofen that improve Purkinje neuron spike regularity without reducing firing frequency. At a disease stage in *Atxn1*^*154Q/2Q*^ mice when motor impairment is due to cerebellar dysfunction, orally administered chlorzoxazone-baclofen improves motor performance without affecting muscle strength.

**Conclusion:** We identify a tight relationship between baclofen-chlorzoxazone concentrations needed to engage target, and levels above which cerebellar function will be compromised. We propose to use this information for a novel clinical trial design, using sequential dose escalation within each subject, to identify dose levels that are likely to improve ataxia symptoms while minimizing toxicity.

## Introduction

The CAG triplet repeat-associated spinocerebellar ataxias (SCA) are a group of neurodegenerative diseases that result in progressive loss of motor function and early death [1]. While the six genetic causes of CAG triplet repeat-associated SCA are unrelated, recent work illustrates that shared underlying mechanisms of disease may contribute to neuronal dysfunction across these disorders [2-5]. SCA1, one of the most common and well-studied of the SCAs, is caused by a glutamine-encoding CAG triplet expansion in the *ATXN1* gene. Cerebellar Purkinje neurons are prominently affected in SCA1 and are believed to underlie motor impairment, while dysfunction of brainstem nuclei and other non-cerebellar structures correlates more closely with early death [6-8]. However, the relative contribution of these structures to motor impairment at various stages of disease remains unclear.

Recent work demonstrates that Purkinje neuron dysfunction is a central cause of motor impairment in rodent models of SCA. In murine models, abnormalities in Purkinje neuron membrane excitability often manifest as irregular, slow, or absent spiking. Interestingly, specific ion-channels have been associated with firing dysfunction across several of these models [2-5], suggesting shared mechanisms of disease and potentially overlapping therapeutic targets. These channels include the large conductance calcium-activated potassium (BK) channel, along with calcium sources that can activate BK function [2]. In the transgenic ATXN1[82Q] mouse model of SCA1, targeting reduced BK channel function with the FDA-approved potassium channel activating compounds chlorzoxazone and baclofen (chlorzoxazone-baclofen) can restore Purkinje neuron spiking and improve motor dysfunction [9]. Due to Purkinje neuron-specific expression of the ATXN1[82Q] transgene in this model of SCA1 [10], the ability of chlorzoxazone-baclofen to improve cerebellar symptoms without worsening non-cerebellar symptoms is not currently known.

The benefit of chlorzoxazone-baclofen on Purkinje neuron spiking has previously been assessed at a single concentration in the transgenic ATXN1[82Q] model of SCA1 [9]. In these mice, Purkinje neurons undergo depolarization block due to profound loss of BK channel current early in disease, causing a majority of cells to cease firing at the onset of symptoms [4]. Since ATXN1[82Q] transgene expression is restricted to Purkinje neurons in this model, it is unclear whether chlorzoxazone-baclofen may have off-target effects outside of the cerebellum. It is therefore imperative to identify the concentration ranges of chlorzoxazone-baclofen that can optimally improve Purkinje neuron function while minimizing extra-cerebellar toxicity. This will enable the design of future clinical trials to assess the safety and efficacy of chlorzoxazone-baclofen in human SCA.

In the present work, we use a genetically precise knock-in mouse model of SCA1 to explore appropriate concentrations and target engagement of chlorzoxazone-baclofen. Our studies demonstrate that BK channels are the likely calcium-activated potassium channel target in SCA1. Importantly, chlorzoxazone-baclofen co-treatment improves motor function in SCA1 mice at an age when the motor phenotype is not confounded by changes in grip strength. In addition, chlorzoxazone-baclofen show no signs of toxicity in SCA1 mice and do not worsen deficits in grip strength at a later disease timepoint when motor weakness is present. These data indicate that chlorzoxazone-baclofen treatment specifically targets cerebellar dysfunction in SCA1, and supports the further development of chlorzoxazone-baclofen as a treatment for cerebellar symptoms in SCA1.

## Methods

### Animals

All animal studies were reviewed and approved by the University of Michigan Institutional Animal Care and Use Committee and were conducted in accordance with the United States Public Health Service’s Policy on Humane Care and Use of Laboratory Animals. *Atxn1*^*154Q/2Q*^ mice, which express an expanded CAG triplet repeat in the endogenous *Atxn1* locus [11], were maintained on a C57BL/6 background. Heterozygous *Atxn1*^*154Q/2Q*^ mice and wild-type littermate controls were used for all studies. All studies were performed at 14 weeks of age or 20 weeks of age. Sexes were balanced for all animal studies.

### Patch clamp electrophysiology

Patch clamp electrophysiology was performed as described previously [5]. Briefly, artificial CSF (aCSF) was prepared in the following manner (in mmol/L): 125 NaCl, 3.8 KCl, 26 NaHCO_3_, 1.25 NaH_2_PO_4_, 2 CaCl_2_, 1 MgCl_2_, 10 glucose. Mice were deeply anesthetized using isoflurane inhalation, decapitated, and brains were rapidly removed for slice preparation. Slices were prepared at 300 µM on a VT1200 vibratome (Leica Biosystems, Buffalo Grove, IL) using the “hot cut” technique [12], and solutions were maintained between 33° and 36°C during tissue sectioning. Slices were sectioned, incubated, and stored in pre-warmed, carbogen-bubbled (95% O_2_, 5% CO_2_) aCSF. For recording, borosilicate glass patch pipets of 3-5 MΩ were filled with an internal pipet solution containing (in mmol/L): 119 K-gluconate, 2 Na-gluconate, 6 NaCl, 2 MgCl_2_, 0.9 EGTA, 10 HEPES, 14 tris-phosphocreatine, 4 MgATP, 0.3 tris-GTP, at pH 7.3 and osmolarity 290 mOsm.

Patch-clamp recordings were performed at 33°C. Pre-warmed, carbogen-bubbled aCSF was perfused at a rate of 150 mL/hour for all recordings. Recordings were acquired using an Axopatch 200B amplifier, Digidata 1440A interface, and pClamp-10 software (Molecular Devices, San Jose, CA). Current clamp data were acquired at 100 kHz in the fast current-clamp mode with bridge balance compensation, and filtered at 2 kHz. Cells were included only if the series resistance did not exceed 15 MΩ at any point during the recording, and if the series resistance did not change by more than 20% during the course of the recording. All presented voltage clamp data have been corrected for a 10 mV liquid junction potential.

### Pharmacology

For some recordings, aCSF contained the following reagents (as specified in the results section for individual experiments): baclofen (Sigma Aldrich, Cat. #5399; 100 nM, 400 nM, or 2 µM), chlorzoxazone (Sigma Aldrich, Cat. #4397; 5 µM, 10 µM, or 50 µM), 6,7-dinitroquinoxaline-2,3-dione (DNQX) (Sigma Aldrich, Cat. #D0540; 10 µM); picrotoxin (Sigma Aldrich, Cat. #P1675; 50 µM). During perfusions of chlorzoxazone and baclofen, a baseline of 5 minutes was established before perfusion of compound. Next, either a single compound or both compounds were perfused for up to twenty minutes. The reported data are taken from the final 2.5 minutes of the baseline recording, and the final 2.5 minutes of the recording in the presence of the perfused compounds. For recordings in the presence of synaptic inhibitors, a 5 minute baseline was similarly established before drug perfusion. DNQX and picrotoxin were then perfused for 5 minutes, at which point the regularity and frequency of firing were compared before and after drug perfusion.

### Tissue Immunofluorescence and Microscopy

Brains from *Atxn1*^*154Q/2Q*^ mice and wild-type littermate controls were placed in 1% paraformaldehyde for 1 hour at room temperature, and were then transferred to a solution of 30% sucrose in phosphate-buffered saline (PBS) for 48 hours at 4°C. Brains were preserved in a 1:1 mixture of 30% sucrose in PBS:OCT compound (Fisher Scientific, Cat. #23-730-571) at - 80°C. Brains were sectioned to 20 µM thickness on a CM1850 cryostat (Leica Biosystems, Buffalo Grove, IL). Tissue was permeabilized with 0.4% triton in PBS, then non-specific binding was minimized by blocking with 5% normal goat serum in 0.1% triton in PBS. Primary antibodies were incubated in PBS containing 0.1% triton and 2% normal goat serum at 4°C overnight.

Primary antibodies used were as follows: mouse anti-BK (1:500; NeuroMab, Cat. #75-022) mouse anti-Ca_V_3.1 (1:150; NeuroMab, Cat. #75-206), rabbit anti-Ca_V_2.1 (1:1000; Abcam, Cat. #ab32642-200), mouse anti-IP3 (1:350; NeuroMab, Cat. #75-035), and mouse anti-Calbindin (1:1000; Sigma Aldrich, Cat. #C9848) or rabbit anti-calbindin (1:1000; Sigma Aldrich, Cat. #C2724) to label Purkinje neurons. Secondary antibody was applied for 1 hour at room temperature in PBS. Secondary antibodies used were as follows: Alexa Fluor 488 goat anti-mouse IgG (H+L) (1:200; Invitrogen, Cat. #A11001), Alexa Fluor 488 goat anti-rabbit IgG (H+L) (1:200; Invitrogen, Cat. #A11034), Alexa Fluor 594 goat anti-rabbit IgG (H+L) (1:200; Invitrogen, Cat. #A11012), Alexa Fluor 594 goat anti-mouse IgG (H+L) (1:200; Invitrogen, Cat. #A11032). Sections were imaged on an Axioskop 2 plus microscope (Zeiss, White Plains, NY) and quantified using ImageJ by measuring mean pixel intensity within a box covering the molecular layer of lobule 5. Representative images were acquired on a C2+ confocal microscope (Nikon, Melville, NY) at 60x magnification.

Cell counts and molecular layer thickness measurements were performed on 20-week-old *Atxn1*^*154Q/2Q*^ mice and wild-type littermate controls after completion of behavioral studies. Tissue was stained for calbindin to mark cerebellar Purkinje neurons, as described above. A distance to 150 µM was drawn from the base of lobule 4, at which point a perpendicular line was drawn from the middle of the soma of the nearest Purkinje neuron to the tip of its dendritic arbor. This distance was recorded as the molecular layer thickness. For cell counts, a distance of 700 µM was measured from the base of lobule 4 and all stained Purkinje neuron somata were counted in that distance. All microscopy and image analyses were performed with the reviewer blind to genotype.

### In Vivo Delivery of Chlorzoxazone-Baclofen

Chlorzoxazone and baclofen were prepared in drinking water as described previously [9]. For longitudinal drug delivery, baclofen (350 µmol/L) and chlorzoxazone (15 mmol/L) were dissolved in drinking water containing 0.05% β-(hydroxypropyl)-cyclodextrin, 40 µL/L NaOH, and 6% sucrose. Vehicle drinking water contained 0.05% β-(hydroxypropyl)-cyclodextrin, 40 µL/L NaOH, and 4% sucrose to ensure that an equal volume of water was consumed across groups. Mice were treated with drinking water beginning at 13 weeks of age. Water was provided *ad libitum* until the end of experimentation, at 20 weeks of age. Water bottles were changed twice weekly.

### Behavior Assays

Motor performance was analyzed using a rotarod protocol as described previously [9]. Mice were handled 3 times before 8 weeks of age to acclimate them to the experimenter. Mice were then trained on the rotarod at 9 weeks of age, using an accelerating speed from 4-40 rpm for 3 consecutive days followed by one day at a constant speed of 24 rpm. At 10 weeks of age, mice were tested for 4 consecutive days at 24 rpm and then randomized into groups based on their baseline performance, keeping baseline consistent between groups within genotype. Drug or vehicle was then administered via water bottles for the duration of the experiment. When the mice reached 14 and 20 weeks of age, they were re-trained for one day and then tested for 4 consecutive days at 24 rpm.

The hanging wire task was performed as described previously [13]. Mice were placed upside down on a suspended wire of 3 millimeter diameter and held approximately 12 inches above a soft landing surface. The time for the mouse to lose grip of the wire was recorded, to a maximum time of 120 seconds. Mice were tested at 14 and 20 weeks of age, on the final day of the rotarod task.

### Statistical Analysis

Data were compiled in Excel (Microsoft Corporation, Redmond, WA) and analyzed using Prism (GraphPad, San Diego, CA). Electrophysiology data were analyzed by using either paired or unpaired Student’s t-test as appropriate. Analyses of grip strength and body weight were completed using a one-way analysis of variance (ANOVA) with a Holm-Sidak correction for multiple comparisons. Analyses of the AHP and rotarod behavioral assays were completed using a two-way repeated measures ANOVA with a Holm-Sidak correction for multiple comparisons. Statistical significance was determined with an α-level of 0.05 for all studies.

## Results

### Purkinje neuron dysfunction is associated with motor impairment preceding neurodegeneration in *Atxn1*^*154Q/2Q*^ mice

Purkinje neuron dysfunction is an early feature of disease in many models of spinocerebellar ataxia (SCA) [3-5, 13-18]. Changes in Purkinje neuron firing are directly relevant to disease, as improving Purkinje neuron firing also improves motor dysfunction in murine SCA models [5, 9, 13, 16-18]. Abnormalities in Purkinje neuron firing are particularly relevant in SCA1, as pharmacologic and genetic strategies to improve Purkinje cell intrinsic excitability improves not just motor dysfunction but also degeneration in murine models of disease [4, 18, 19]. Prior studies in SCA1 have focused primarily on the transgenic ATXN1[82Q] model of SCA1, where polyglutamine expanded ATXN1 expression is restricted to Purkinje cells. While the ATXN1[82Q] model of SCA1 accurately models cerebellar features of disease, the relevance of cerebellar dysfunction to the overall motor phenotype is unknown in the more genetically precise *Atxn1*^*154Q/2Q*^ model of SCA1, in which a polyglutamine-expanded *Atxn1* transgene is knocked into the endogenous *Atxn1* locus, and therefore widely expressed in the nervous system and elsewhere. In this model of SCA1 with a hyperexpanded polyglutamine repeat motor dysfunction and premature death have been attributed, at least in part to spinal cord and brainstem motor neuron degeneration [7, 11]. We therefore sought to identify whether cerebellar dysfunction is present, and relevant to the motor phenotype in *Atxn1*^*154Q/2Q*^ mice.

We examined Purkinje neuron membrane excitability in *Atxn1*^*154Q/2Q*^ mice at 14 weeks of age, an age at which motor impairment is prominent [11]. No evidence of neurodegeneration is present in *Atxn1*^*154Q/2Q*^ mice even at 20 weeks of age (Supplementary Figure 1A-B). When compared to Purkinje neurons from wild-type littermate controls, *Atxn1*^*154Q/2Q*^ Purkinje neurons display irregular spiking (Figure 1A-C) with no change in firing frequency (Figure 1D). Since Purkinje neuron firing in SCA murine models differs between the anterior and posterior cerebellum [2, 5], we independently analyzed neurons from the anterior cerebellum (lobules IV-V) and nodular zone (lobule X) and found a similar increase in spike irregularity with no change in firing frequency across both cerebellar regions. Additionally, since alterations in Purkinje neuron physiology may reflect changes in either intrinsic pacemaking or synaptic activity, we performed recordings in *Atxn1*^*154Q/2Q*^ and wild-type control Purkinje neurons before and after the addition of synaptic inhibitors (6,7-dinitroquinoxaline-2,3-dione [DNQX], 10 µM; picrotoxin, 50 µM). No significant difference in either spike frequency (Supplementary Figure 1C) or regularity (Supplementary Figure 1D) was noted after the addition of synaptic inhibitors, suggesting that altered Purkinje neuron spiking in *Atxn1*^*154Q/2Q*^ mice is intrinsically driven. When neurons were held at -80 mV and injected with depolarizing current, we found that *Atxn1*^*154Q/2Q*^ neurons require a significantly lower amplitude of injected current to elicit depolarization block than wild-type littermate control neurons (Figure 1E). Importantly, these alterations in spiking were accompanied by no significant change in input resistance between *Atxn1*^*154Q/2Q*^ and wild-type littermate control neurons (Figure 1F).

**Figure 1.**
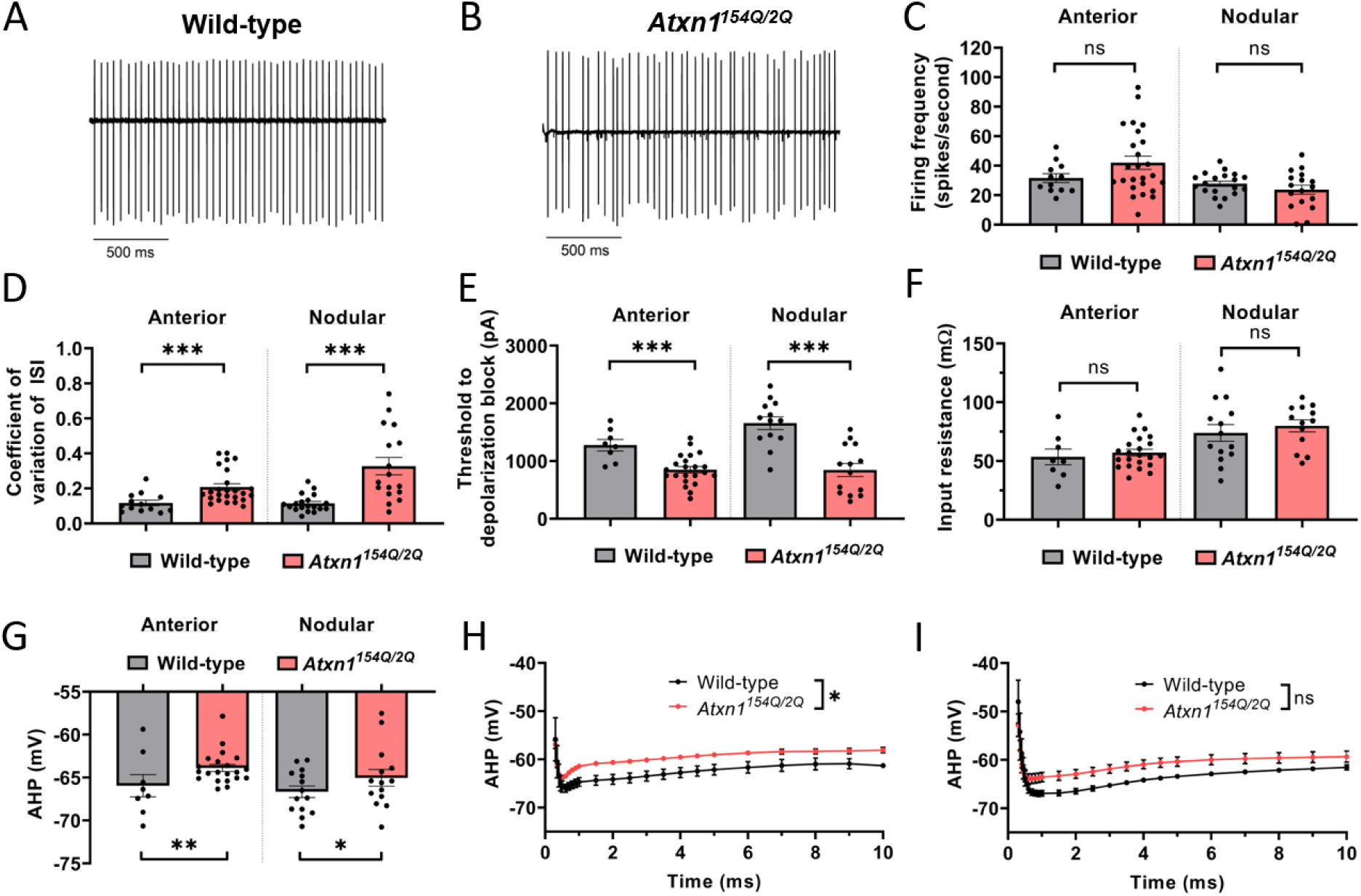
Alterations in Purkinje neuron physiology in *Atxn1*^*154Q/2Q*^ mice. Representative spontaneous Purkinje neuron spiking is shown for 14 week-old (A) wild-type and *(B) Atxn1*^*154Q/2Q*^ mice. (C) Firing frequency in wild-type and *Atxn1*^*154Q/2Q*^ Purkinje neurons. (D) Regularity of Purkinje neuron spiking, as indicated by the coefficient of variation of the interspike interval, in wild-type and *Atxn1*^*154Q/2Q*^ mice. (E) Threshold of injected current required to elicit depolarization block in wild-type and *Atxn1*^*154Q/2Q*^ mice. (F) Input resistance in wild-type and *Atxn1*^*154Q/2Q*^ Purkinje neurons. (G) Minimum spike amplitude, indicative of the fast afterhyperpolarization (AHP) in wild-type and *Atxn1*^*154Q/2Q*^ Purkinje neurons. Quantification of the AHP and interspike interval of wild-type and *Atxn1*^*154Q/2Q*^ Purkinje neurons in the (H) anterior cerebellar lobules and (I) posterior cerebellar lobules. * denotes p<0.05; ** denotes p<0.01, *** denotes p<0.001; ns denotes p>0.05. Two-tailed Student’s t-test (C-G); two-way repeated measures ANOVA with Holm-Sidak correction for multiple comparisons (H-I).

Since irregular Purkinje neuron spiking is strongly associated with deficits in the afterhyperpolarization (AHP) in various mouse models of ataxia [3-5, 20], we examined differences in spike waveform between *Atxn1*^*154Q/2Q*^ Purkinje neurons and wild-type littermate controls. We found that the AHP in *Atxn1*^*154Q/2Q*^ mice was significantly depolarized when compared to wild-type littermate control cells (Figure 1G-I). No other significant changes in spike waveform were noted in *Atxn1*^*154Q/2Q*^ Purkinje neurons when compared to wild-type littermate controls (Table 1). Together, these data indicate that a specific reduction in AHP amplitude accompanies irregular spiking and neuronal hyperexcitability in *Atxn1*^*154Q/2Q*^ mice at a stage of disease with prominent motor impairment.

**Table 1.** Electrophysiological parameters of Purkinje neurons from 14 week-old *Atxn1*^*154Q/2Q*^ mice and wild-type littermate controls. Data are presented as mean ± standard error of the mean. Two-tailed Student’s t-test.

|  | WT<br>anterior | SCA1<br>anterior | P value | WT<br>posterior | SCA1<br>posterior | P value |
| --- | --- | --- | --- | --- | --- | --- |
| **Firing frequency** | 31.65 ±<br>2.96 | 42.04 ±<br>4.47 | 0.1359 | 27.81 ±<br>1.76 | 23.78 ±<br>3.12 | 0.255 |
| **Coefficient of variation**<br>**of ISI** | 0.1171 ±<br>0.015 | 0.2082<br>± 0.019 | **<0.001** | 0.1145 ±<br>0.011 | 0.3274 ±<br>0.050 | **<0.001** |
| **AHP minimum** | -65.93 ±<br>1.30 | -63.88 ±<br>0.415 | **0.0073** | -66.65 ±<br>0.676 | -65.03 ±<br>0.967 | **0.0259** |
| **Spike peak** | 15.37 ±<br>1.83 | 14.17 ±<br>1.13 | 0.5882 | 21.27 ±<br>0.82 | 18.52 ±<br>1.88 | 0.216 |
| **Depolarization dV/dt** | 479.1 ±<br>17.5 | 472.5 ±<br>16.9 | 0.8296 | 574.6 ±<br>21.5 | 513.4 ±<br>26.9 | 0.0936 |
| **Repolarization dV/dt** | -359.9 ±<br>22.5 | -352.3 ±<br>9.93 | 0.7237 | -336.4 ±<br>27.5 | -343.5 ±<br>20.1 | 0.8347 |
| **Spike half-width** | 0.245 ±<br>0.016 | 0.241 ±<br>0.005 | 0.7446 | 0.273 ±<br>0.021 | 0.2492 ±<br>0.013 | 0.3231 |
| **Input resistance** | 53.51 ±<br>6.66 | 57.21 ±<br>2.85 | 0.5529 | 73.97 ±<br>7.11 | 79.96 ±<br>5.11 | 0.5058 |

Changes in ion channel expression or function accompany changes in Purkinje neuron spiking in mouse models of SCA [14]. Additionally, the observed irregular spiking and reduced AHP amplitude observed in *Atxn1*^*154Q/2Q*^ Purkinje neurons is consistent with calcium-activated potassium (K_Ca_) channel dysfunction or voltage-gated calcium (Ca_V_) channel dysfunction [4, 16, 20-22]. We examined previously published microarray data to identify potential ion channel transcript dysregulation in *Atxn1*^*154Q/2Q*^ and identified reduced transcripts for *Kcnma1, Cacna1g, Trpc3* and *Itpr1* [2, 23]. Reduced *Kcnma1* and *Cacna1g* transcripts were observed at 14 weeks of age in *Atxn1*^*154Q/2Q*^ mice when compared to wild-type littermate controls [2]. We therefore sought to quantify protein expression of large conductance K_Ca_ (BK) channels (encoded by *Kcnma1*) and putative calcium sources that may activate BK, including Ca_V_3.1 (*Cacna1g*) [5], Ca_V_2.1 (*Cacna1a*) [24], and the IP3 receptor (*Itpr1*) [25] in *Atxn1*^*154Q/2Q*^ mice and wild-type littermate controls at 14 weeks of age. In both the anterior cerebellum (Figure 2A) and nodular zone (Figure 2B), fluorescence intensity of BK and Ca_V_3.1 (Figure 2C) was reduced in the cerebellar molecular layer of *Atxn1*^*154Q/2Q*^ mice compared to wild-type littermate controls, while the IP3 receptor and Ca_V_2.1 (Figure 2C) showed no change in expression. Together, these data suggest that reduced cerebellar expression of BK and Ca_V_3.1 underlies the altered physiology observed in *Atxn1*^*154Q/2Q*^ mice.

**Figure 2.**
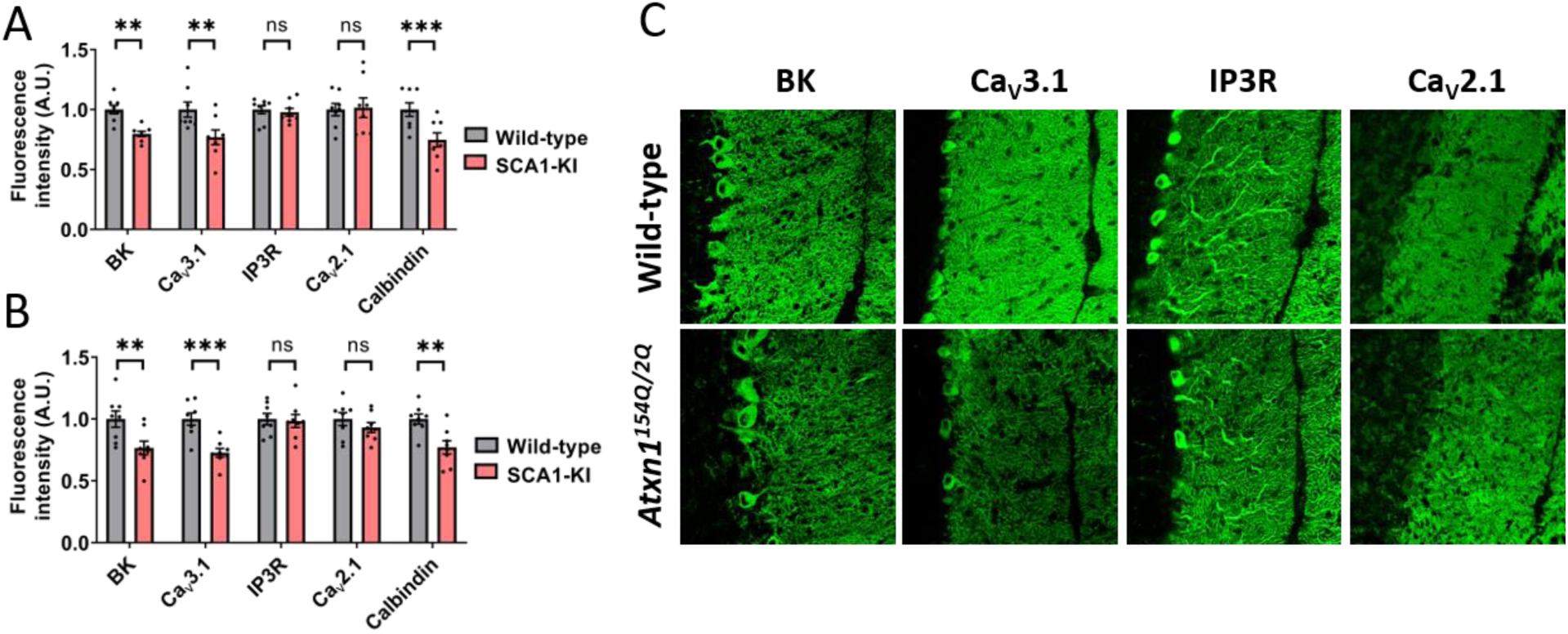
Reduced expression of BK and Ca_V_3.1 channels in *Atxn1*^*154Q/2Q*^ cerebellum. (A) Ion channel expression was measured in the anterior cerebellum (lobule 5) molecular layer of 14 week-old *Atxn1*^*154Q/2Q*^ mice and wild-type littermate controls. (B) Ion channel expression was measured in the nodular zone (lobule 10) molecular layer of 14 week-old *Atxn1*^*154Q/2Q*^ mice and wild-type littermate controls. (C) Representative 40x confocal microscopy images are shown for (C) BK, Ca_V_3.1, IP3R, and Ca_V_2.1 in *Atxn1*^*154Q/2Q*^ mice and wild-type littermate controls. ** denotes p<0.01; *** denotes p<0.001; ns denotes p>0.05. Two-tailed Student’s t-test (F), with Holm-Sidak correction for multiple comparisons used in (A-B).

### A combination of chlorzoxazone and baclofen improves *Atxn1*^*154Q/2Q*^ Purkinje cell firing

The altered Purkinje neuron spiking observed in symptomatic *Atxn1*^*154Q/2Q*^ mice is similar to neuronal dysfunction seen in another mouse model of SCA1, the ATXN1[82Q] transgenic model [4]. The more profound loss of BK and Cav3.1 channel expression in ATXN1[82Q] mice results in a more prominent electrophysiologic phenotype, in which neurons undergo a complete loss of spiking [4], which nevertheless represents a continuum with the irregular spiking observed in *Atxn1*^*154Q/2Q*^ Purkinje neurons [2] (Figure 1). The shared underlying dysfunction of BK channels and reduced AHP amplitude suggests that BK channels may be an important target in SCA1.

In transgenic ATXN1[82Q] mice, combined treatment with chlorzoxazone and baclofen targets potassium channel dysfunction to improve Purkinje neuron spiking and motor impairment at both early- and mid-symptomatic ages [9]. Chlorzoxazone is a canonical K_Ca_ channel activator that can target BK channels [26]. Baclofen, a GABA_B_ receptor agonist, is a known activator of subthreshold-activated potassium channels in Purkinje neurons [27]. Subthreshold-activated potassium channels can compensate for loss of the AHP and facilitate repetitive spiking in Purkinje cells [3]. In ATXN1[82Q] mice, a combination of chlorzoxazone and baclofen (chlorzoxazone-baclofen) was necessary to achieve adequate K_Ca_ channel activation and restoration of function [9]. We wished to determine the necessity of a combination of these agents to improve the more modestly impaired firing in *Atxn1*^*154Q/2Q*^ mice. Since SCA1-KI mice exhibit a gradation of Purkinje cell firing abnormalities, this allows for investigation of dose-dependence of chlorzoxazone-baclofen to correct irregular spiking. It is important to determine the optimal concentrations of chlorzoxazone-baclofen as excessive activation of K^+^ channels results in reduction in Purkinje cell firing frequency, which is independently correlated with motor dysfunction (reviewed in [28]).

The effect of different doses of chlorzoxazone and baclofen were assessed in acute cerebellar slices from 14-week *Atxn1*^*154Q/2Q*^ mice by perfusing varying doses of either individual compounds or combinations of chlorzoxazone-baclofen. A high combined dose of chlorzoxazone (50 µM) and baclofen (2 µM) was able to improve spike regularity in *Atxn1*^*154Q/2Q*^ Purkinje neurons, as indicated by a reduction in the coefficient of variation (CV) of the interspike interval (ISI) (Figure 3A). An intermediate combined dose of chlorzoxazone (10 µM) and baclofen (400 nM) similarly reduced the CV of the ISI in *Atxn1*^*154Q/2Q*^ neurons (Figure 3A), while a lower combined dose of chlorzoxazone (5 µM) and baclofen (100 nM) had no effect on CV of the ISI (Figure 3A). Firing frequency was examined in these same recordings. Firing frequency was greatly suppressed by the high combined dose of chlorzoxazone (50 µM) and baclofen (2 µM), while no effect on firing frequency was observed at the intermediate combined dose of chlorzoxazone (10 µM) and baclofen (400 nM) (Figure 3B). A small reduction in firing frequency was observed in the low combined dose of chlorzoxazone (5 µM) and baclofen (100 nM) (Figure 3B). Interestingly, when treated with the intermediate dose of chlorzoxazone alone (10 µM; Figure 3C) or baclofen alone (400 nM; Figure 3D), the CV of the ISI was not changed from baseline. This is contrast to perfusion of higher dose chlorzoxazone alone (50 µM) or baclofen alone (2 µM), which both reduce the CV of the ISI (Supplemental Figure 2A-B) but also suppress firing frequency (Supplemental Figure 2C-D). Taken together, these data suggest that chlorzoxazone-baclofen improves Purkinje neuron spiking in a dose-responsive manner in *Atxn1*^*154Q/2Q*^ mice, and that the intermediate dose of a combination of chlorzoxazone (10 µM) and baclofen (400 nM) has the optimal effect of improving Purkinje neuron spiking irregularity without affecting firing frequency in acute cerebellar slices. Additionally, these data clearly indicate the necessity of a combination of chlorzoxazone and baclofen to achieve the effect of improving spike regularity while not affecting firing frequency.

**Figure 3.**
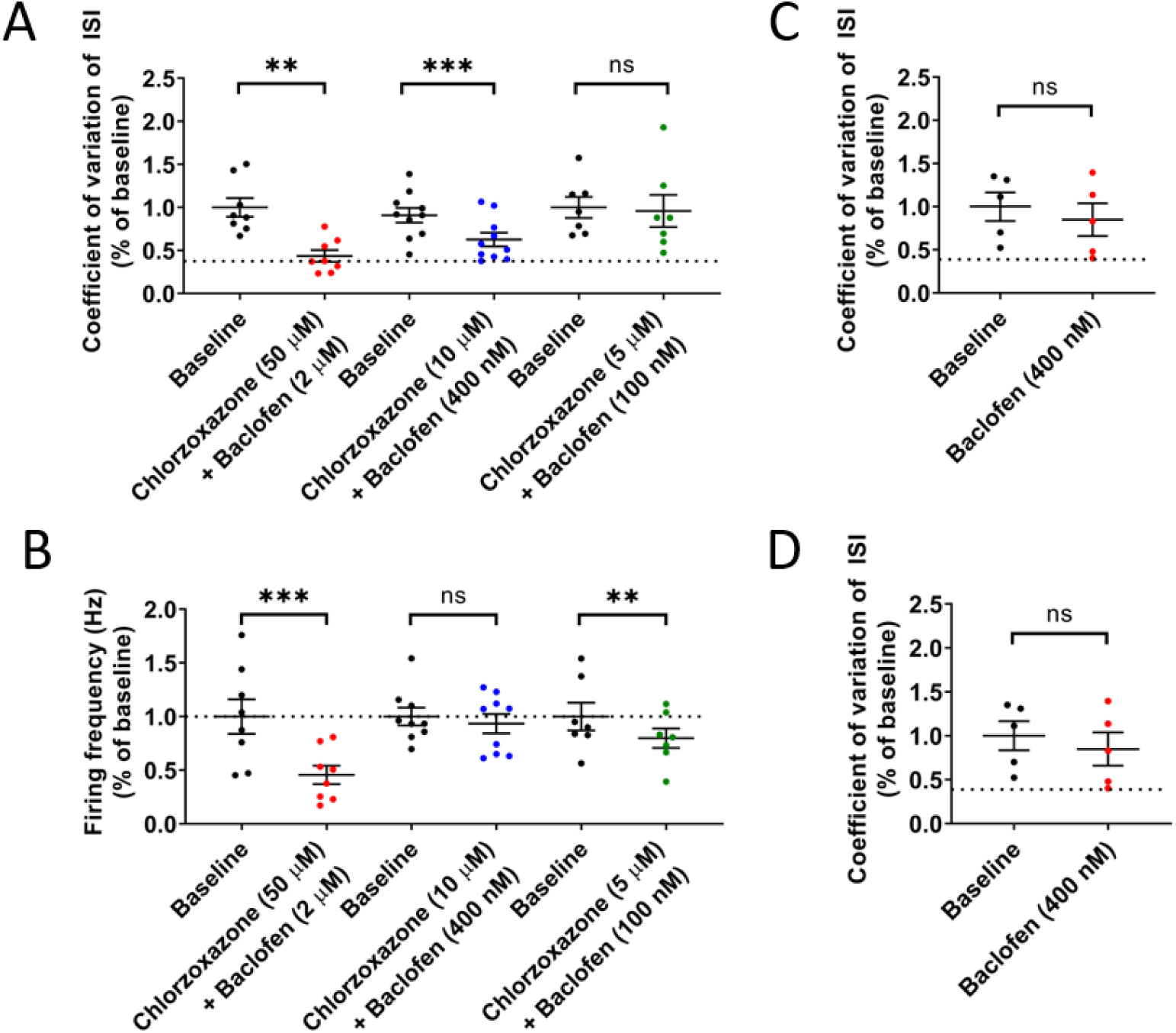
Chlorzoxazone-baclofen improves *Atxn1*^*154Q/2Q*^ Purkinje neuron physiology in a dose-responsive manner. Acute cerebellar slices were prepared from 14 week-old *Atxn1*^*154Q/2Q*^ mice for all recordings. (A) Regularity of Purkinje neuron spiking was recorded by measuring the coefficient of variation (CV) of the interspike interval (ISI). CV of the ISI was recorded at baseline and normalized to 1.0, and was then recorded after perfusion of differing concentrations of chlorzoxazone and baclofen into the bath. Dotted line represents normalized CV of the ISI from age-matched wild-type littermate controls. (B) Firing frequency was recorded at baseline and normalized to 1.0, and was then recorded after perfusion of differing concentrations of chlorzoxazone and baclofen into the bath. Dotted line represents normalized baseline firing frequency from age-matched wild-type littermate controls. (C) Similar to (A), CV of the ISI was recorded at baseline and after perfusion of an intermediate dose of chlorzoxazone (10 µM). Dotted line represents normalized CV of the ISI from age-matched wild-type littermate controls. (D) Similar to (C), CV of the ISI was recorded at baseline and after perfusion an intermediate dose of baclofen (400 nM). Dotted line represents normalized CV of the ISI from age-matched wild-type littermate controls. * denotes p<0.05; ** denotes p<0.01; *** denotes p<0.001; ns denotes p>0.05. Paired Student’s t-test.

### Chlorzoxazone and baclofen specifically target cerebellar dysfunction in SCA1

*Atxn1*^*154Q/2Q*^ mice have a hyperexpanded polyglutamine repeat. In human SCA1, large repeat expansions are associated with brainstem predominant disease and premature death in the second decade of life [29-31]. It is important to know whether cerebellar dysfunction contributes to motor impairment in *Atxn1*^*154Q/2Q*^ mice and accurately serves as a model for testing therapy for cerebellar dysfunction, the major source of disability and quality of life in human SCA1.

*Atxn1*^*154Q/2Q*^ mice are described to have motor neuron involvement [7], which may confound interpretation of the rotarod task in mice. We therefore evaluated grip strength, a measure of motor neuron function, using the hanging wire task. At 14 weeks of age, no impairments in grip strength were noted in *Atxn1*^*154Q/2Q*^ mice when compared to wild-type littermate controls (Figure 4A). *Atxn1*^*154Q/2Q*^ mice were administered chlorzoxazone-baclofen in drinking water at a dose expected to achieve the intermediate concentrations (see Figure 4 of [9]) of chlorzoxazone (1-10 µM) and baclofen (100 nM-1 µM) in the brain. Importantly, chlorzoxazone-baclofen did not affect grip strength of 14 week-old *Atxn1*^*154Q/2Q*^ mice (Figure 4A). The rotarod assay is commonly used to test balance and gait coordination, and reflects balance impairment when not confounded by impaired muscle strength [32]. At 14 weeks, *Atxn1*^*154Q/2Q*^ mice display significantly impaired motor function on the rotarod assay when compared to wild-type littermate controls (Figure 4B). The absence of grip-strength impairment in the *Atxn1*^*154Q/2Q*^ mice provides more confidence that the rotarod impairment at this stage of disease is due to cerebellar dysfunction. Treatment with chlorzoxazone-baclofen significantly improved motor performance in 14 week-old *Atxn1*^*154Q/2Q*^ mice (Figure 4B).

**Figure 4.**
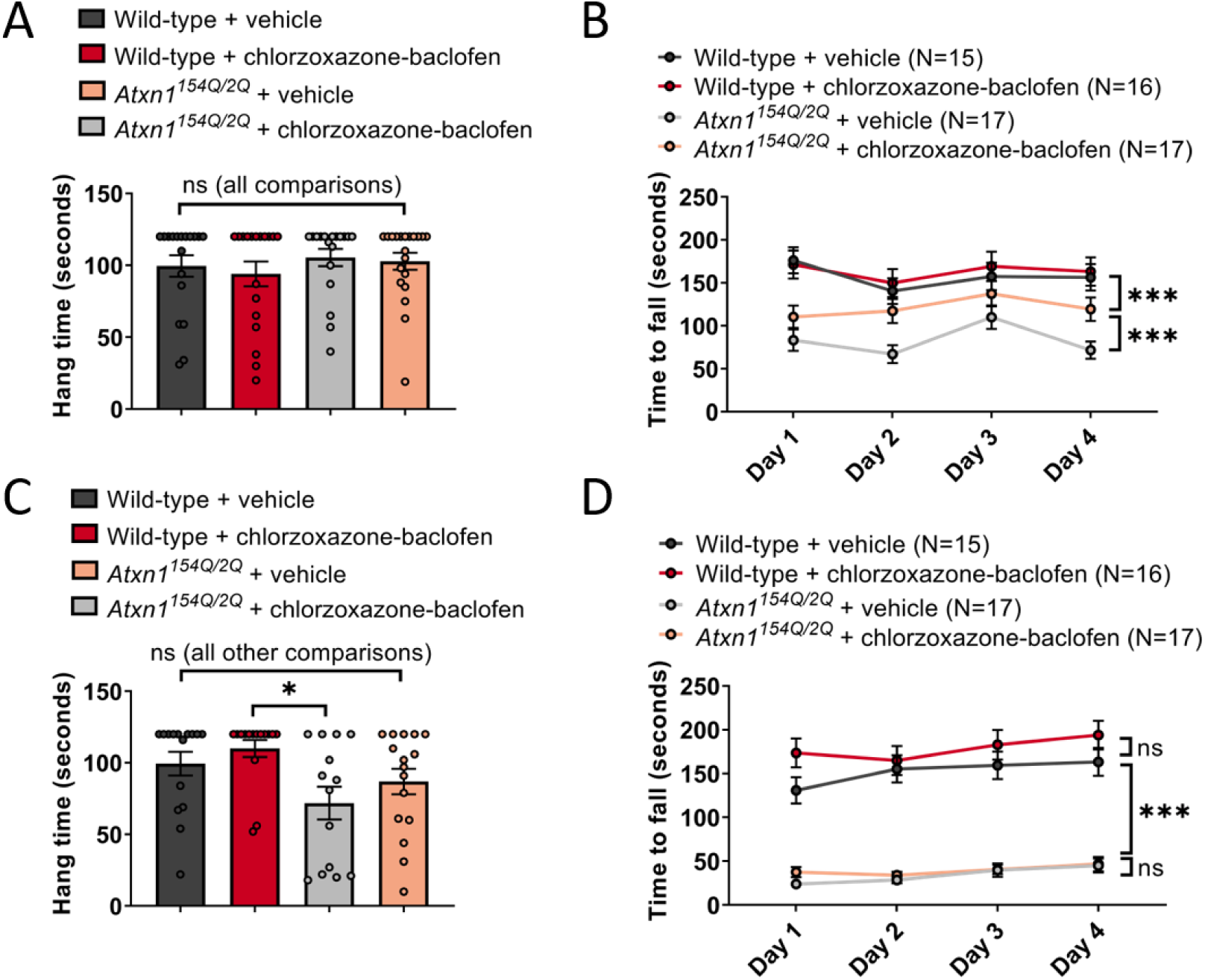
Chlorzoxazone-baclofen improves cerebellar motor impairment in *Atxn1*^*154Q/2Q*^ mice. *Atxn1*^*154Q/2Q*^ mice and wild-type littermate controls were treated with chlorzoxazone-baclofen in water bottles starting at 13 weeks of age. (A) Grip strength was measured in *Atxn1*^*154Q/2Q*^ mice and wild-type littermate controls at 14 weeks of age using a hanging wire task. (B) The time to fall off of a rotarod at constant speed was recorded for *Atxn1*^*154Q/2Q*^ mice and wild-type littermate controls at 14 weeks of age. (C) Similar to (A), at 20 weeks of age. (D) Similar to (B), at 20 weeks of age. * denotes p<0.05; ** denotes p<0.01; *** denotes p<0.001; ns denotes p>0.05. Two-way repeated measures ANOVA with Holm-Sidak correction for multiple comparisons (A, C); one-way ANOVA with Holm-Sidak correction for multiple comparisons (B, D).

At 20 weeks of age, a later stage of disease, *Atxn1*^*154Q/2Q*^ mice display a significant impairment in grip strength compared to wild-type littermate controls (Figure 4C). Reassuringly, chlorzoxazone-baclofen, which are FDA approved skeletal muscle relaxants, did not worsen the grip strength deficits in *Atxn1*^*154Q/2Q*^ mice (Figure 4C). At this stage of disease, it would be expected that rotarod performance of *Atxn1*^*154Q/2Q*^ mice would be further impaired, referable to the observed grip strength deficits. Consistent with this hypothesis, 20-week *Atxn1*^*154Q/2Q*^ mice displayed significantly impaired motor performance on the rotarod compared to wild-type littermate controls to a larger degree than was observed at 14 weeks of age. Chlorzoxazone-baclofen did not rescue the rotarod impairment in *Atxn1*^*154Q/2Q*^ mice at this stage of disease (Figure 4D), but, importantly, also did not worsen motor performance in 20 week-old *Atxn1*^*154Q/2Q*^ mice, consistent with the lack of effect of chlorzoxazone-baclofen on grip strength at this age.

Since chlorzoxazone-baclofen is being considered for clinical use in patients with ataxia [9], we evaluated for potential toxicity of long-term use of this combination in *Atxn1*^*154Q/2Q*^ mice. As reported previously, *Atxn1*^*154Q/2Q*^ mice fail to gain weight compared to wild-type littermate controls, which is evident in both male and female mice at both 14 (Figure 5A-B) and 20 (Figure 5C-D) weeks of age [11]. Chlorzoxazone-baclofen did not accelerate this failure to gain weight. A slightly lower baseline weight was observed in chlorzoxazone-baclofen treated wild-type mice (Figure 5A), which was also noted in these same mice at 20 weeks of age (Figure 5C).

**Figure 5.**
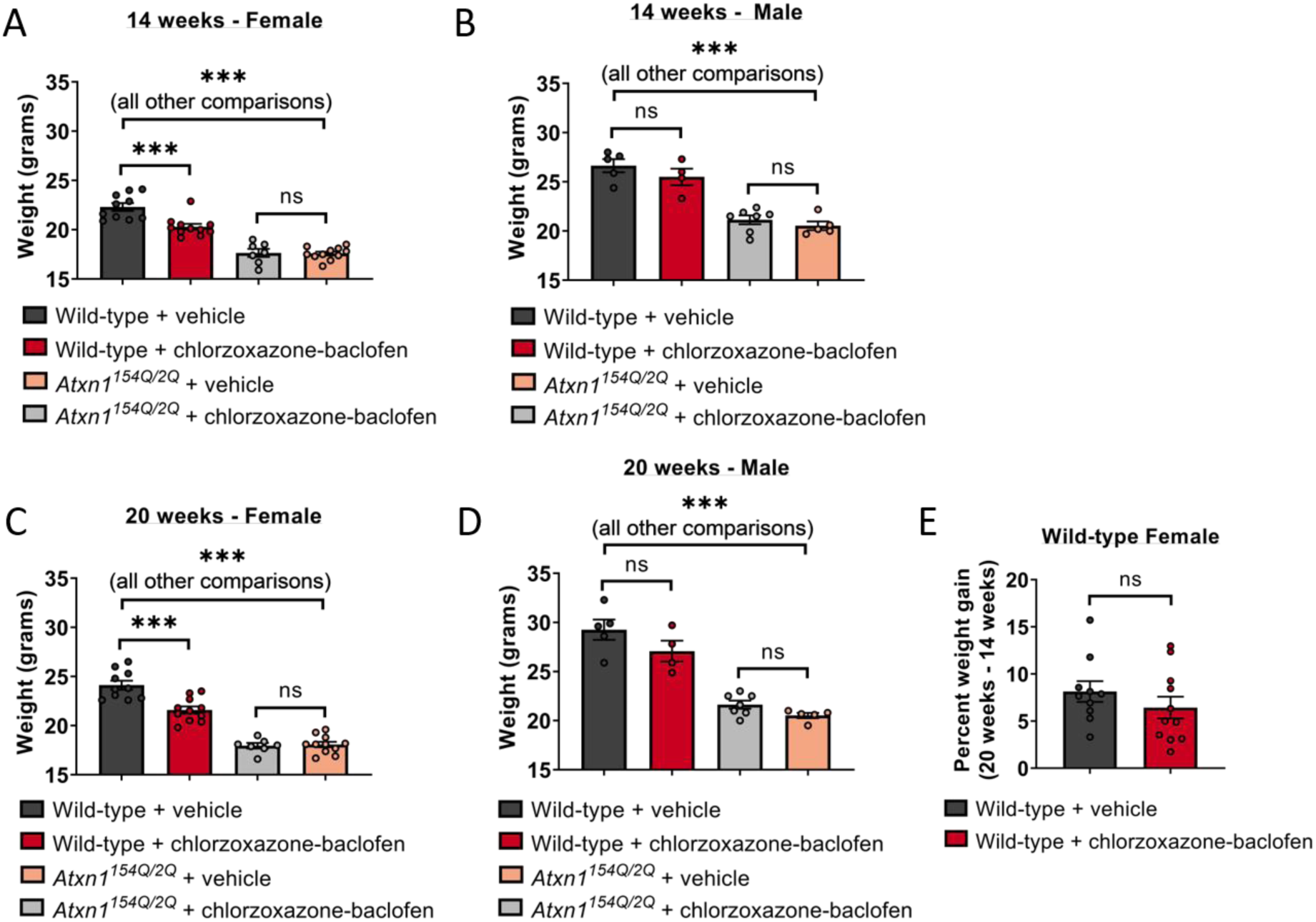
Body weight is preserved in *Atxn1*^*154Q/2Q*^ mice treated with chlorzoxazone-baclofen. Body weight was measured in 14 week-old (A) male and (B) female *Atxn1*^*154Q/2Q*^ mice and wild-type littermate controls after one week of water bottle treatment of chlorzoxazone-baclofen. Similar to (A-B), body weight was measured in 20 week-old (C) male and (D) female *Atxn1*^*154Q/2Q*^ mice and wild-type littermate controls after 7 weeks of continuous chlorzoxazone-baclofen water bottle treatment. (E) Percent change in body weight from 14 to 20 weeks of age was calculated for wild-type female mice treated with either vehicle drinking water or water containing chlorzoxazone-baclofen. *** denotes p<0.001; ns denotes p>0.05. One-way ANOVA with Holm-Sidak correction for multiple comparisons (A-D); Two-tailed Student’s t-test (E).

However, this difference in weight was not accelerated in chlorzoxazone-baclofen treated female wild-type mice when compared to vehicle treated controls at the two recorded timepoints (Figure 5E), suggesting that these mice still gain weight normally and are not experiencing toxicity from chlorzoxazone-baclofen treatment. Together, these behavioral studies indicate that at 14 weeks, cerebellar dysfunction contributes meaningfully to motor impairment in *Atxn1*^*154Q/2Q*^ mice and can be ameliorated by a treatment strategy targeting cerebellar dysfunction. At a later disease stage, *Atxn1*^*154Q/2Q*^ mice display a motor phenotype that is driven by impairments of muscle strength. These findings reinforce the idea that chlorzoxazone-baclofen specifically targets motor impairment due to cerebellar dysfunction in SCA1.

## Discussion

Chlorzoxazone-baclofen has been proposed for the treatment of cerebellar motor dysfunction in SCA [9]. However, the optimal dose of chlorzoxazone-baclofen for target engagement without off-target toxicity is not known. In the present study, we identify a dose range of chlorzoxazone-baclofen that improves Purkinje neuron spike regularity in the *Atxn1*^*154Q/2Q*^ model of SCA1 without suppressing firing frequency. Importantly, chlorzoxazone-baclofen improves cerebellar motor dysfunction in *Atxn1*^*154Q/2Q*^ mice at an age when motor impairment is cerebellar in origin, and does not negatively affect grip strength. At a later stage of disease, when motor impairment is worsened by impaired grip strength, chlorzoxazone-baclofen does not cause further impairment in motor function. Taken together, these studies suggest that chlorzoxazone-baclofen can be dosed to properly engage target in the cerebellum while simultaneously minimizing off-target toxicity. Human clinical trials of chlorzoxazone-baclofen will determine whether this drug combination may be effective to also treat cerebellar motor impairment in human SCA.

In the transgenic ATXN1[82Q] mouse model of SCA1, chlorzoxazone-baclofen improves cerebellar Purkinje neuron physiology and motor impairment [9]. In the present study, chlorzoxazone-baclofen similarly improves motor function in *Atxn1*^*154Q/2Q*^ mice at an age when motor dysfunction is cerebellar in origin. In ATXN1[82Q] mice, the benefit of chlorzoxazone-baclofen persists at later stage disease, despite prominent Purkinje neuron dendritic degeneration seen at this disease stage [9]. Chlorzoxazone-baclofen was, however, unable to sustain improved motor function in *Atxn1*^*154Q/2Q*^ mice in later stage disease. This discrepancy may be explained by the genetic differences in these two models. The transgenic ATXN1[82Q] model of SCA1 overexpresses mutant *ATXN1* specifically in cerebellar Purkinje neurons under control of the *Pcp2* promoter [10]. Conversely, *Atxn1*^*154Q/2Q*^ mice express glutamine-expanded *Atxn1* at its endogenous locus, resulting in widespread expression throughout the central nervous system [11]. As a result, motor impairment in ATXN1[82Q] mice is cerebellar in origin, even in late stage disease, and largely reflects Purkinje neuron dysfunction. Spinal motor neuron involvement in *Atxn1*^*154Q/2Q*^ mice likely contributes to motor impairment in this model of SCA1 [7, 11]. The current study suggests that motor dysfunction in mid-stage disease in *Atxn1*^*154Q/2Q*^ mice is driven by cerebellar dysfunction, as evidenced by the improvement in motor function with chlorzoxazone-baclofen, whereas later stage motor impairment is a result of motor neuron dysfunction that is not improved by chlorzoxazone-baclofen. In this regard, *Atxn1*^*154Q/2Q*^ mice likely recapitulate features of human SCA1, where morbidity is primarily cerebellar in origin in earlier stages of disease, whereas end stage disease is due to cranial motor neuron dysfunction. Chlorzoxazone and baclofen are used clinically as muscle relaxants and are not recommended for simultaneous dosing in patients due to concerns about tolerability [33]. A major concern of using central muscle relaxants in SCA is the potential to worsen motor impairment [34].

However, the present data suggest that chlorzoxazone-baclofen can engage target in cerebellar Purkinje neurons while avoiding toxicity resulting from impaired muscle strength. An optimal effect of combined chlorzoxazone and baclofen treatment would be to improve the regularity of spiking in SCA1 Purkinje neurons without slowing firing frequency, as both irregular spiking and slow Purkinje neuron firing are associated with motor impairment in mouse models of ataxia [5, 14-17, 20, 35]. In the current study, we identify a tight relationship between drug concentrations needed to engage target, and levels above which cerebellar function will be compromised. In planning for a human clinical trial with this drug combination to improve cerebellar ataxia, it will be important to identify the specific orally administered dose of the combination of chlorzoxazone-baclofen that achieves appropriate brain levels. We propose that a scheme of sequential dose escalation within each clinical trial-subject would be the most effective way to identify doses of chlorzoxazone-baclofen that engage target optimally without compromising cerebellar function, by allowing for monitoring of plasma and potentially cerebrospinal fluid levels of chlorzoxazone-baclofen along with effects on cerebellar motor function. In the current study, a high cerebellar concentration of chlorzoxazone-baclofen (50 µM and 2 µM, respectively) impairs cerebellar function, while a low concentration of chlorzoxazone-baclofen (5 µM and 0.2 µM, respectively) fails to engage target. An intermediate concentration of chlorzoxazone-baclofen (10 µM and 0.4 µM, respectively) at the site of action in the cerebellum is associated with optimal improvements in cerebellar physiology. Brain concentrations of baclofen are ∼5 fold lower than plasma concentrations in both humans and mice [36]. Our studies in mice suggest that brain and plasma concentrations of chlorzoxazone reach near equilibrium in mice following chronic administration [9]. We therefore propose, based on the *in vitro* data obtained in the current study, that a plasma concentration range between 8-40 µM chlorzoxazone and 1-4 µM baclofen will be associated with improvement in cerebellar function in human SCA. It is likely that the relationship between administered drug dose, plasma/CSF concentrations of chlorzoxazone-baclofen, and changes in cerebellar function with each drug dose will need to be closely monitored for individual trial participants in a potential efficacy study.

Interestingly, the ion channels that underlie altered Purkinje neuron physiology in *Atxn1*^*154Q/2Q*^ mice are also implicated in neuronal dysfunction in other mouse models of SCA. These channels, BK (encoded by *Kcnma1*) and Ca_V_3.1 (encoded by *Cacna1g*), show reduced expression in another mouse model of SCA1 [2], several mouse models of SCA2 [2], and a mouse model of SCA7 [5]. BK channels rely upon intracellular calcium for activation, and recent evidence suggests that Ca_V_3.1 may be an important calcium source for BK [2, 5]. Despite SCA1, SCA2, and SCA7 resulting from polyglutamine expansions in distinct genes, dysfunction of a BK channel functional module appears to contribute to altered Purkinje neuron spiking and cerebellar motor impairment in all of these disorders [2]. As illustrated in the present study, and in the ATXN1[82Q] model of SCA1 [9], chlorzoxazone-baclofen improves Purkinje neuron physiology through engagement of potassium channel targets. Therefore, it is possible that chlorzoxazone-baclofen may have relevance for treating motor impairment beyond SCA1. This possibility has not been tested in rodent models, although a related genetic strategy of viral BK channel replacement improves Purkinje neuron physiology in SCA7 [5]. It is tempting to speculate that a shared mechanism of dysfunction exists across multiple etiologies of SCA, and that a treatment strategy such as chlorzoxazone-baclofen may therefore have wider relevance for targeting cerebellar motor impairment.

## Author Roles

1. Research project: A. Conception, B. Organization, C. Execution; 2. Statistical Analysis: A. Design, B. Execution, C. Review and Critique; 3. Manuscript Preparation: A. Writing of the first draft, B. Review and Critique

DDB 1A, 1B, 1C, 2A, 2B, 3A, 3B

HH 1B, 1C

LJM 1C

LMM 1C, 2B, 3B

VGS 1A, 1B, 2C, 3A, 3B

---

**Supplementary Figure 1.**
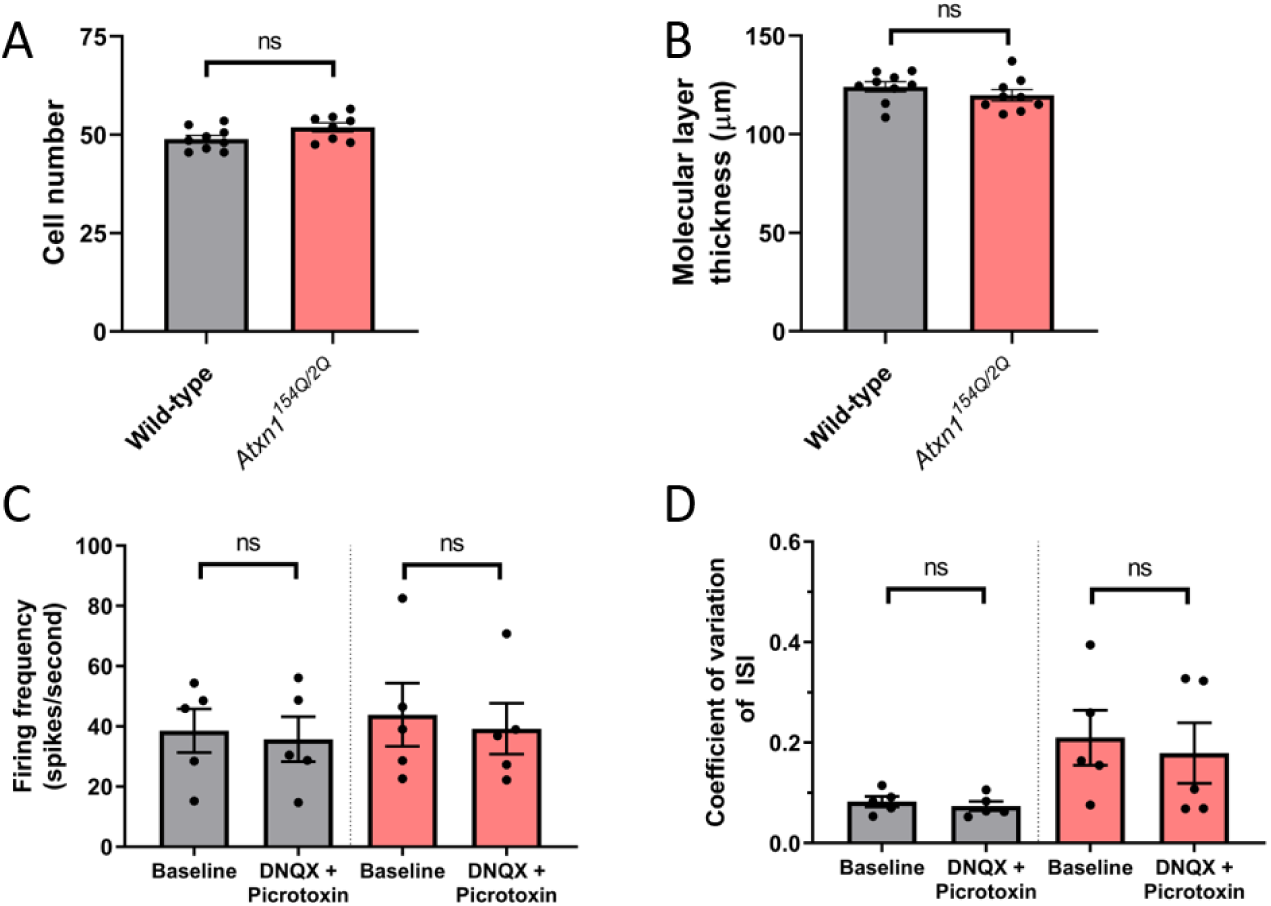
Neurodegeneration and Electrophysiology in *Atxn1*^*154Q/2Q*^ Purkinje neurons. (A) Purkinje neuron cell number in lobule 5 (anterior cerebellum) of wild-type and *Atxn1*^*154Q/2Q*^ mice at 20 weeks of age. (B) Purkinje neuron dendritic length, as indicated by thickness of the cerebellar molecular layer in lobule 5 of wild-type and *Atxn1*^*154Q/2Q*^ mice at 20 weeks of age. (C) Purkinje neuron firing frequency in the anterior cerebellum of 14 week-old wild-type (left, grey) and *Atxn1*^*154Q/2Q*^ (right, red) mice before and after the addition of the synaptic inhibitors 6,7-dinitroquinoxaline-2,3-dione [DNQX] (10 µM) and picrotoxin (50 µM). (D) Similar to (C), indicating spike regularity by measuring the coefficient of variation of the interspike interval (ISI). ns denotes p>0.05. Two-tailed Student’s t-test (A-B), paired Student’s t-test (C-D).

**Supplementary Figure 2.**
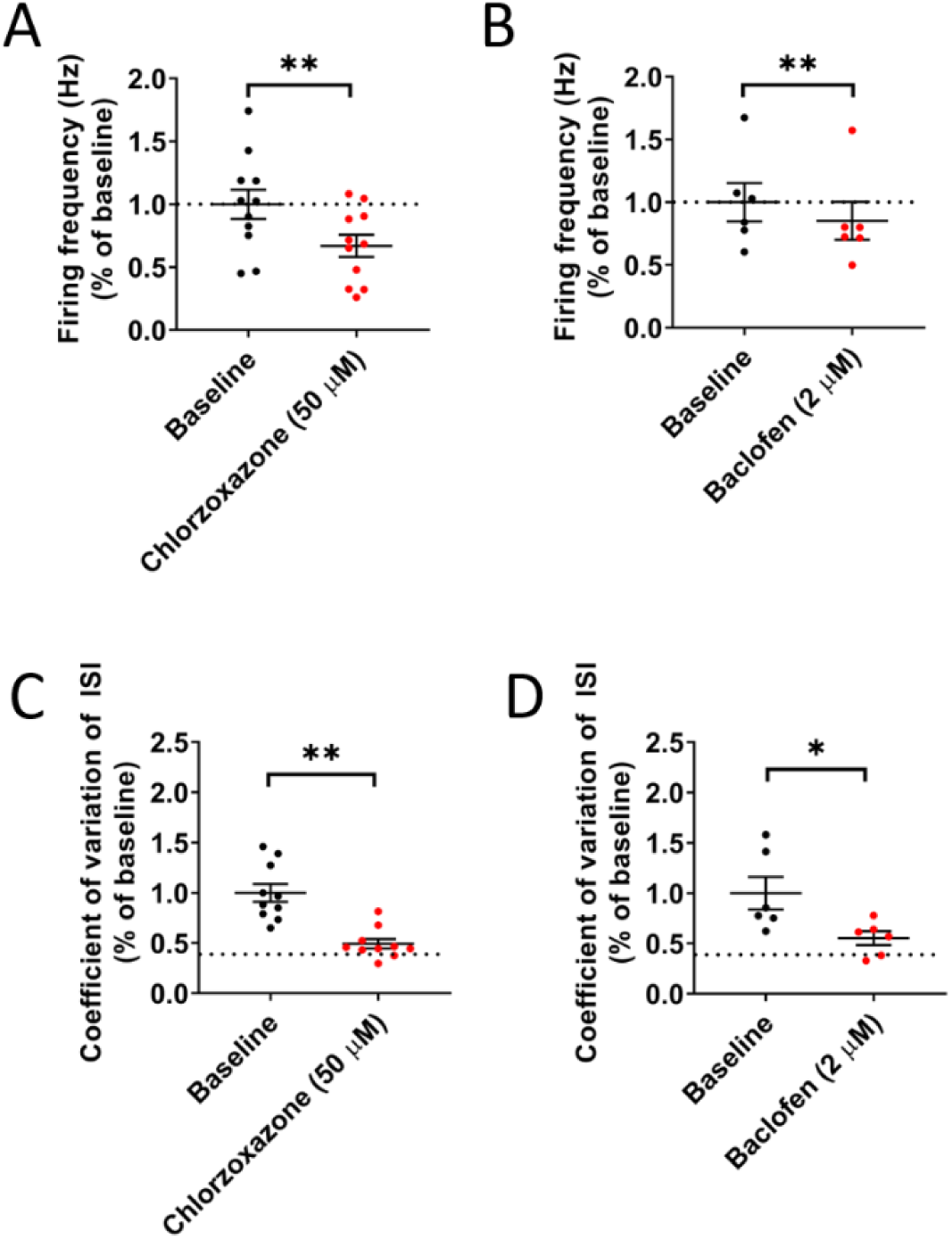
Effect of high-dose chlorzoxazone alone and baclofen alone on Purkinje neuron spiking in *Atxn1*^*154Q/2Q*^ mice. Acute cerebellar slices from 14 week-old *Atxn1*^*154Q/2Q*^ mice were prepared, and Purkinje neurons from the anterior cerebellar lobules were used for all studies. Firing frequency and CV data are normalized to the baseline values of each individual neuron. (A) Measurements of firing frequency before and after perfusion of high-dose chlorzoxazone (50 µM). (B) Measurements of firing frequency before and after perfusion of high-dose baclofen (2 µM). (C) Measurements of CV before and after perfusion of high-dose chlorzoxazone (50 µM). (D) Measurements of CV before and after perfusion of high-dose baclofen (2 µM). * denotes p<0.05; ** denotes p<0.01. Paired Student’s t-test.

